# Low-dimensional factorized neural computations underlie risk-adaptive choices

**DOI:** 10.64898/2026.08.12.744223

**Authors:** T Alexander Price, Andrew Liu, Rhiannon L Cowan, Niloufar Shahdoust, Tyler S Davis, Bornali Kundu, John D Rolston, Shervin Rahimpour, Ben Shofty, Alla Borisyuk, Elliot H Smith

## Abstract

Real-world decision-making rarely occurs with perfect information. Instead, individuals must constantly weigh potential rewards against the probability of adverse outcomes.^1^ Failures of this process can lead to maladaptive decisions associated with reduced lifetime success, and numerous psychiatric disorders such as gambling addictions, bulimia nervosa, and substance use disorder.^2,3^ The neural computations that facilitate inference about the landscape of potential outcomes remain unclear, but are thought to occur in distributed frontotemporal circuits.^4^ Here we used deep reinforcement learning agents to predict distinct behavioral strategies and their underlying neural population dynamics during a risky decision-making task. Across a range of training conditions, deep reinforcement learning agents separated into strategies marked by either overly cautious exploration of the reward contingency space or a high-performing, risk-adaptive Bimodal strategy. The internal dynamics of high-performing Bimodal agents formed low-dimensional representations that segregated safe and risky states. In contrast, the cautious exploration agents were associated with more skewed and entangled neural representations. We found remarkably similar dynamical representations and their associated behavioral strategies in neuronal ensemble recordings from human epilepsy patients performing a similar risky decision-making task. These results reveal the structure of dynamical computations that underlie inferences about uncertain outcomes and their associated behavioral strategies.

## Main

Reinforcement learning (RL) provides a powerful framework for modeling how agents navigate environments to maximize their expected reward.^5^ Advancements in artificial intelligence have provided several solutions for improving agents’ performance in uncertain environments, including contextualizing the uncertainty of environmental states and their probabilistic transitions,^6^ or providing information about the full distribution of potential rewards.^7^ The latter strategy has been discovered in biological neural networks, allowing organisms to learn from distributions of abstract and temporally delayed rewards as a result of efficient coding in midbrain dopamine neurons.^8–11^ While many RL algorithms aim to maximize expected value of the reward, their learning rules can give rise to qualitatively distinct policies, just as biological agents frequently demonstrate distinct preferences in uncertain environments, such as risk-aversion or risk-seeking.^12^ These risk-related traits can be associated with depression, impulsivity, or adverse, maladaptive outcomes across the lifespan.^3,13^

Recent theoretical advances have shown that neural computations often occur on low-dimensional manifolds: low-dimensional geometric structures in neuronal ensemble activity space that constrain and define the possible computations an ensemble can perform.^14–19^ Applying this dynamical systems perspective to human decision-making opens avenues for comparing biological and artificial intelligence. If deep RL agents and human brains converge on similar behavioral strategies to solve complex tasks, do they also converge on similar internal representations? Recent work in “neuro-AI” (artificial intelligence) has begun to explore these parallels, using the internal state dynamics of artificial agents to generate hypotheses about biological neural manifolds.^20–22^ Specifically, the geometry of artificial neural population dynamics may reveal similarities in how risk and uncertainty are managed by human neural networks.^23^

We bridged these domains by using deep RL agents to generate hypotheses about the behavioral strategies and neural representations in humans performing an externally valid risky decision-making task called the Balloon Analog Risk Task (BART).^24–28^ Deep RL agents settled on distinct behavioral phenotypes that emerged spontaneously in both artificial agents and human participants: an efficient, risk-adaptive ‘Bimodal’ strategy and a cautious, less successful ‘Explorer’ strategy. By analyzing the dynamics of the deep RL agents’ recurrent layer activity, we discovered neural trajectories evolving around a dominant axis that represented expected outcomes. Remarkably, we found that humans adopting a Bimodal behavioral strategy exhibited the same geometric neuronal population structure as their artificial counterparts. These findings suggest that specific dynamical computations support model-based inference and efficient risk management in uncertain environments.

### Learning the internal dynamical structure of risky choices with deep RL agents

In order to generate predictions about the diversity of neural computations and algorithms for solving BART, a total of 400 randomly initialized deep RL agents with the Actor-Critic architecture (Fig. 1**a**) were trained on a version of the BART called meta-BART (Fig. 1**b**, Supp. Fig. 1**a**).^28,29^ During each trial in the BART (or meta-BART for the agents; see Methods) the goal is to earn points by inflating a balloon whose radius signifies potential risk and reward; larger balloons are worth more points, popped balloons earn no reward. During training on the meta-BART, epochs of 50 balloons had their maximum sizes drawn from a normal distribution around a random mean, *µ* ([*µ*s between 0.2 and 1]; Fig. 1b). Agents were trained until they met a performance threshold and their network weights were subsequently saved. On average this took about 330,000 time steps (2.74 checkpoints). These agents then completed an evaluation suite of meta-BART with 17 preselected balloon size averages. The activity of the nodes in the recurrent layer for each of the agents was recorded and saved for analysis.

**Figure 1.**
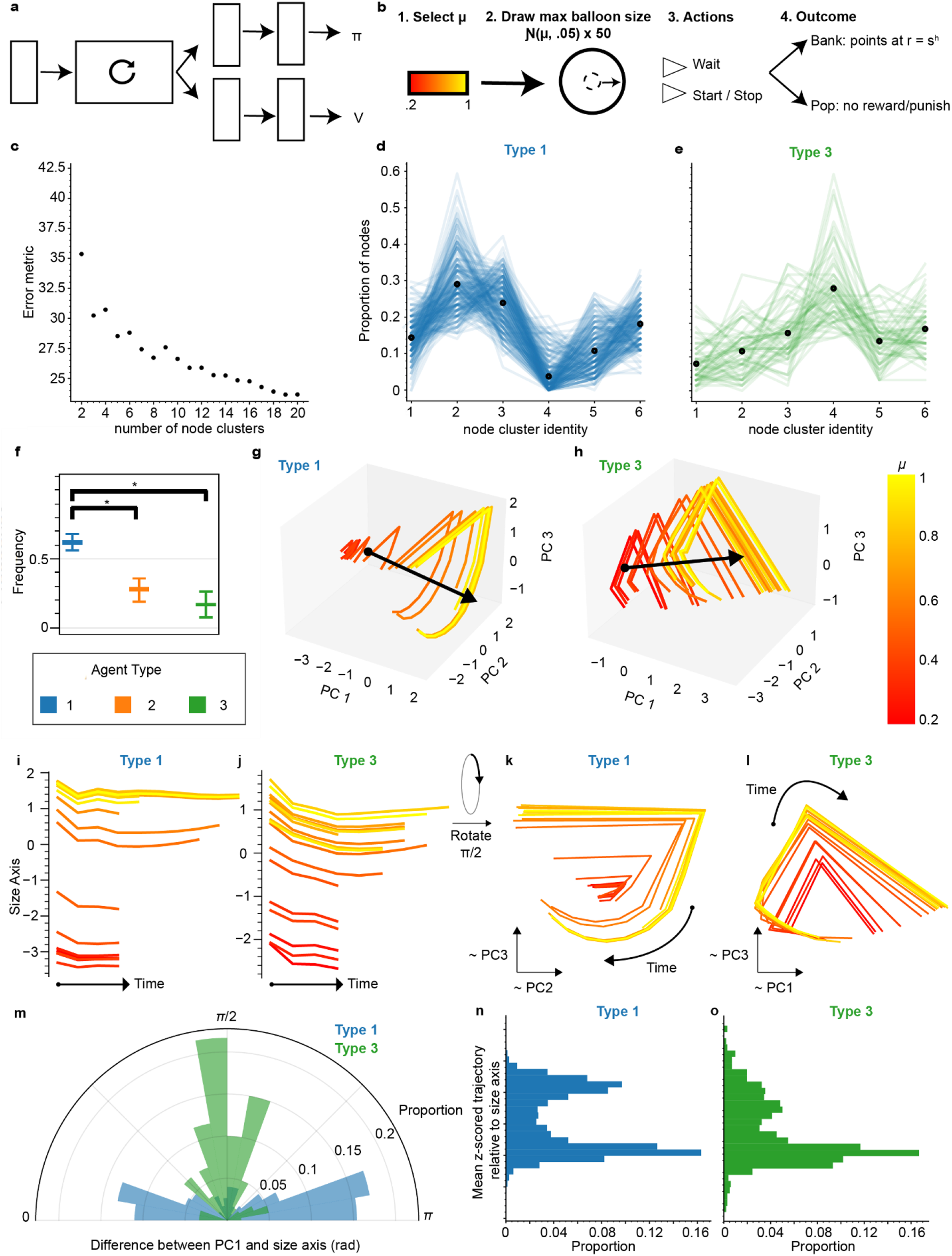
Varied strategies and neural dynamics underlie risk and reward estimates in deep RL agents. **a**, Actor-critic architecture of the deep RL agents with each box representing a feed-forward layer (64-nodes) and the circular arrow representing the recurrent layer (64-nodes). **b,** Schematic of the meta-BART task. (1) During a 50-balloon epoch, a mean balloon size, µ, was drawn (training) or selected (testing) from a uniform distribution. (2) Each individual balloon size was then drawn from a normal distribution, N(µ,0.05). (3) At each time step, agents could take one of two actions, either start/stop the inflation, or allow the balloon to continue to inflate. (4) Finally, if the agent ceased inflation they were rewarded based on the size of the balloon. In some cases, agents were punished for popping a balloon. **c,** k-means clustering of the activity of nodes in the recurrent layers of all agents. K=12 was selected as the elbow, which then reduced to 6 clusters when the repeated and inverted activities were removed. **d,e,** Agent types based on proportion of node types in recurrent layer. Each line represents an individual agent and black dots are means for each cluster type for each agent type. **f,** Proportions of each agent type (n_Type 1_ = 227, n_Type 2_ = 113, n_Type 3_ = 60). **g,h,** Representative trajectories of node activities of Type 1 (**g**) and Type 3 (**h**) agent example plotted in 3D PCA space. The black arrow represents a decoded size axis projected into the same PCA space. **i,j,** The PCA trajectories projected onto the size axis for both the Type 1 (**i**) and Type 3 (**j**) agent. **k,l,** View along the size axis for Type 1 (**k**) and Type 3 (**l**) agents. **m,** Polar histogram of cosine angles between PC1 and size axis for Type 1 (blue) and Type 3 (green) agents. **n,o,** Histograms of the mean value along the size axis for each µ for (**n**) Type 1 and (**o**) Type 3 agents.

We clustered the activity patterns of nodes across all agents’ recurrent layers using k-means clustering (k = 12; Fig. 1**c**, See Methods). This procedure grouped nodes with similar patterns of activity over the course of the whole meta-BART session (Supp. Fig. 1**b**). All cluster centers had some amount of near-periodic behavior following the natural evolution of the meta-BART task as balloons were inflated, popped or banked, and then reset to the next balloon with a size of zero. Increasing the number of clusters did not generate cluster centers with new features in the dynamics. We then classified agents into three groups based on the proportion of their node types (k-means, k=3; Fig. 1**d,e**). About 56.8% of the agents were classified as Type 1 agents and about 15.0% were classified as Type 3 agents (Fig. 1**f**). Interestingly, Type 1 agents reached the performance threshold after an average of 2.5 checkpoints while Type 3 agents took 4.2 checkpoints on average. Type 2 agents did not have a clear, identifiable behavioral approach and were thus no longer considered (Supp. Fig 1). These agents seemed to be a mix of Type 1 and Type 3 agents from their distribution of nodes.

To investigate the dynamical computations carried out by deep RL agents, we plotted neural trajectories comprising the top 3 principal components (PCs) of recurrent layer activations for each agent (Fig. 1**g,h**). We noticed that neural trajectories evolved as parallel arcs distributed along a linear axis encoding balloon size. We sought to understand how the extent of the arcs around the balloon size axis related to agents’ balloon size inferences by estimating the convex hulls (akin to surface area) of the arcs. For smaller balloons (*µ* < 0.55), convex hulls were significantly smaller in Type 1 agents than in Type 3 agents (Supp. Fig. 2), suggesting a discrepancy in agents’ representations of smaller and larger balloons.

**Figure 2.**
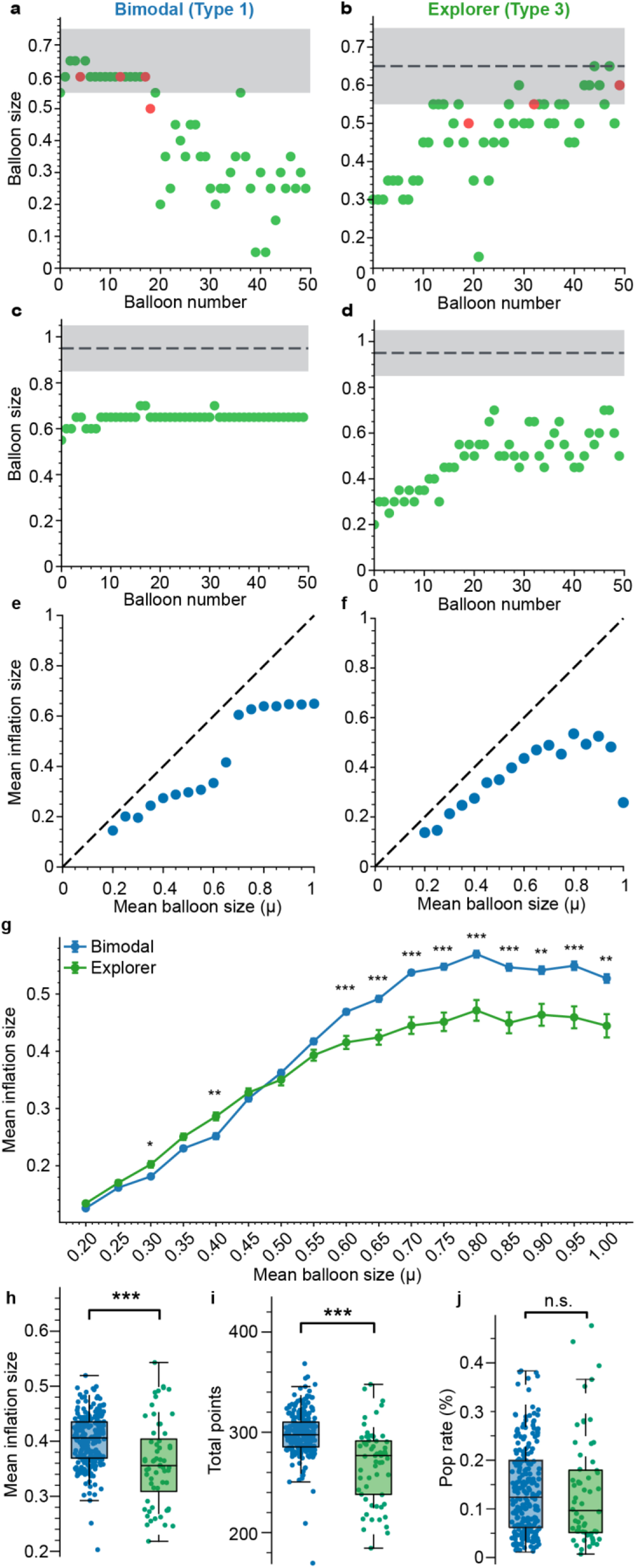
Distinct classes of risk-adaptive behavior in deep RL agents. **a-d**, Different single 50-balloon epochs for an example Type 1 (Bimodal; **a,c**) and Type 3 (Explorer; **b,d**) agents. **a,b,** µ = 0.65, **c,d,** µ = 0.95. Green circles indicate rewarded balloons, red circles indicate popped balloons. The black dotted line represents the selected µ, and the shaded gray area is two standard deviations from the selected mean. **e-f,** Mean inflated balloon size for the same Bimodal (**e**) and Explorer (**f**) agent (as **a,c** and **b,d** respectively) across all µs presented in the evaluation suite. **g,** Mean inflation sizes on the selected µs in the evaluation suite across each class of agent. **h-j,** Box and scatter plots of mean Inflation Size, total scores and pop rates of the population of agents. Blue data is from Bimodal agents, green data is from Explorer agents. Error bars in panel **g** are STD. *, **, *** indicate statistical significance (p<0.05, p<0.01, p<0.001). N_Bimodal_ = 227, N_Exploer_ = 60.

We next sought to understand how balloons of various sizes were oriented along the balloon size axis, hypothesizing that the axis modeled agents’ inferred balloon sizes. We estimated the balloon size axis via a partial least squares regression, orienting it such that larger balloons aligned in the positive direction. The neural trajectories for Type 1 agents clustered bimodally, near the edges of the size axis (Fig. 1**i,k**), whereas Type 3 agent exhibited unimodal, small-skewed distributions of trajectories along the balloon size axis (Fig. 1**j,l**). Both Types 1 and 3 agents exhibited balloon size axes, however, the balloon size axis in Type 1 agents aligned with PC1 (cosine angle 27.7 ± 25.2 degrees), whereas the balloon size axis for Type 3 agents was organized orthogonally (cosine angle 69.3 ± 25.0 degrees; Fig. 1**m**). We verified that these trajectory distribution shapes were indicative of the full populations of Type 1 and Type 3 agents by plotting histograms of their z-scored mean trajectories (Fig. 1**n,o**). Aggregated Type 1 agents had a significant Bimodal distribution of mean size projections while the Type 3 agents did not (*Hartigan and Hartigan Dip Tests; Type 1: p < 0.001, Type 3: p = 0.47*). In the following text, we thus refer to these agent types as Bimodal (Type 1) and Explorer (Type 3), respectively.

### Distinct classes of risk-adaptive behavior arise from agents’ neural classifications

After clustering agents by their node activity and characterizing their dynamical representations of expected reward (via maximizing balloon size), we sought to understand whether these classes of learned internal representations distinguished their task behavior. Originally classified via the proportion of node types in their recurrent layers, agents’ behavior on the evaluation suite divided neatly into distinct behavioral classes (*χ*^2^*(4) = 58.59, p < 0.05*). Type 3 agents acted as cautious Explorers, using the full 50-balloon epoch to slowly discover each balloon size mean (Fig. 2**b,d**), whereas Type 1 agents did not frequently explore the full balloon size space (Fig. 2**a,c**). Instead, Type 1 agents favored a more efficient and effective Bimodal inflation time (IT) strategy, where they were more likely to select a lower mean IT for smaller *µ* and a higher mean IT for larger *µ* (Fig. 2**e**) compared to Type 3 agents (Fig. 2**f**).

When the mean ITs of the agentic populations were compared, the Explorer population inflated smaller balloon sizes (*µ* = 0.2 to *µ* = 0.45) further, including at two *µ*s where their mean inflation size was significantly larger than the Bimodal population’s mean inflation size (two-tailed t-test with Bonferroni correction for multiple comparisons, *µ = 0.3, t = −3.24, p_adj_ = 0.027 and µ = 0.4, t= −4.03, p_adj_ = 0.0016*, respectively; Fig. 2**g**). However, for larger balloon sizes, the Bimodal population had significantly larger mean inflation sizes, from *µ* = 0.6 to *µ* = 1 (two tailed t-test with Bonferroni correction for multiple comparisons; Fig. 2**g**). As a result of differences in behavioral strategy there were significant differences in overall task success. Bimodal agents both inflated balloons significantly longer overall (*two-tailed t-test, t= 5.11, p < 0.001*; Fig. 2**h**) and scored significantly more points across the evaluation suite (*two-tailed t-test, t = 7.99, p < 0.001*; Fig. 2**i**), without exhibiting a significantly higher pop rate *(two-tailed t-test, t = 0.32, p = 0.75* Fig. 2**j**).

### Deep RL agent categories predict risk-adaptive behavioral strategies in humans

After observing the distinct strategies that the agents developed from random initialization, and their underlying internal representations, we sought to test whether these strategies predicted human participants’ performance during BART. To accomplish this, 44 patients (23 female) with drug-resistant epilepsy who were undergoing neuromonitoring completed 47 sessions of a computerized BART (three patients completed the task twice; number of trials = 239.2 ± 21.3). In this version of BART, participants inflated one of three colors of balloons drawn from three corresponding size distributions, and therefore potential rewards (see representative trial in Fig. 3**a**, top). Red balloons popped at the smallest average size, orange balloons popped at an intermediate size, and yellow balloons popped at the largest size.

**Figure 3.**
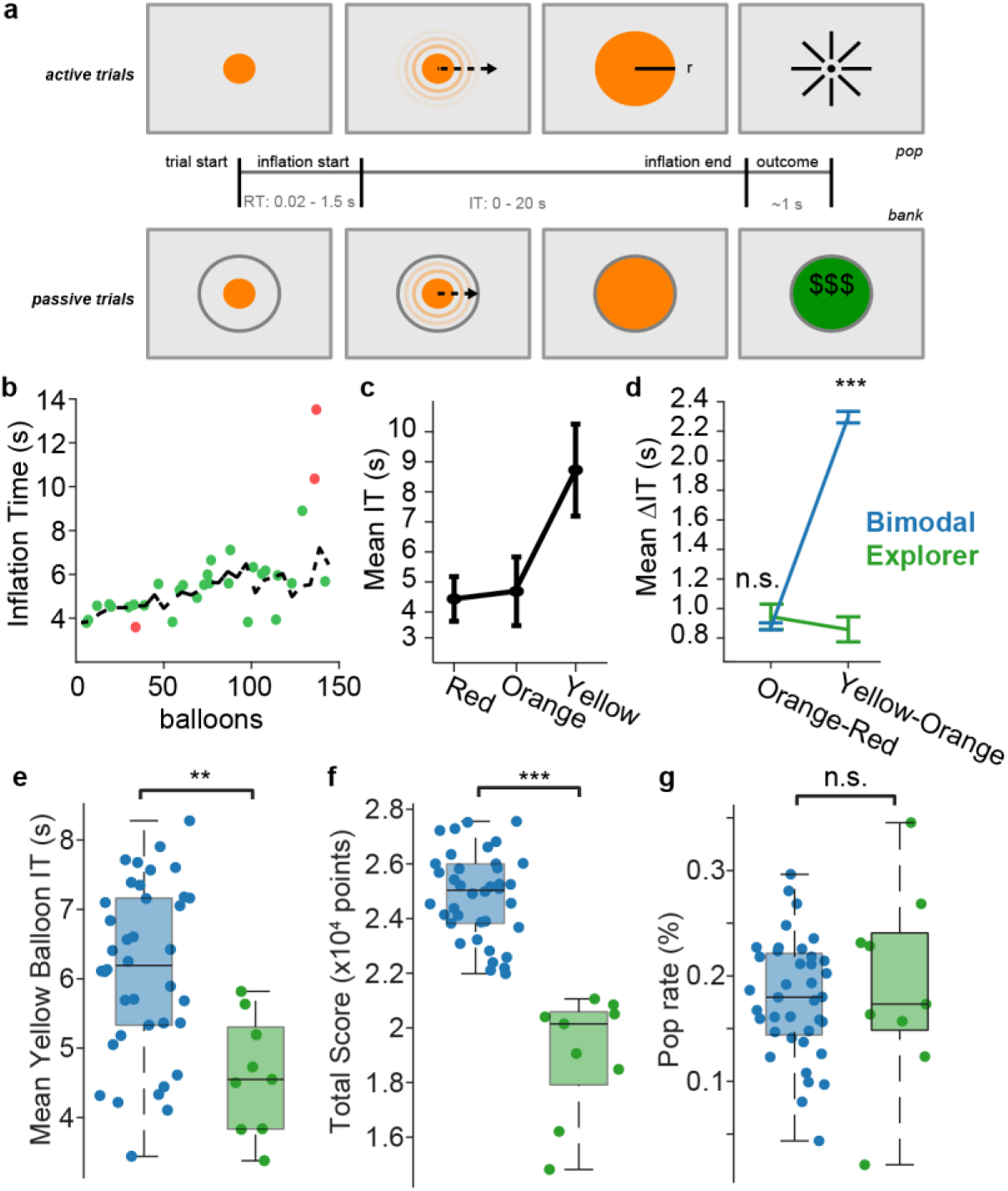
Deep RL agents predict human behavioral strategies. **a**, Schematic of human version of continuous BART. **b,** An example of a human participant using an Explorer strategy on yellow balloons. **c,** An example of a human participant using a Bimodal strategy. **d,** An analysis of human subpopulation ITs showing differences between the mean IT of orange balloons and the mean IT of red balloons and differences between the mean ITs of yellow balloons and the mean ITs of orange balloons. **e-g,** Human participants using a Bimodal strategy had significantly longer ITs (**e**) and higher score (**f**). Pop rates were also not significantly different between groups of humans (**g**). Error bars in panels **c,d** are STD. *, **, *** indicate statistical significance (p<0.05, p<0.01, p<0.001). N_Bimodal_ = 38, N_Explorer_ = 9

We used k-means clustering to split the participants into behavioral groups based on several task-derived behavioral measures (i.e., total score, pop rate and mean ITs for red, orange and yellow balloons; Supp. Fig. 2). Participants’ sessions were clustered into two groups whose proportions corresponded with the two categories of deep RL agents: Bimodal (n = 38) and Explorer groups (n = 9; Supp Fig. 3). We noticed the behaviors of these groups were similar to the two classes of agents, with Explorer participants slowly increasing their ITs over the course of the task (Fig. 3**b**), and Bimodal participants learning to leverage increased potential rewards of yellow balloons (Fig. 3**c**). To verify the correspondence between the human behavioral groups and behavioral strategies predicted by the agents, we carried out several further tests.

First, mean differences in ITs among the three balloon colors were consistent with the original definition of a ‘Bimodal’ strategy, derived from the agents (*two-tailed t-test, Orange-Red: t = 0.32, p = 0.75; Orange-Yellow: t = −3.72, p = 5.4E-4;* Fig. 3**d**). Moreover, the behavioral strategies developed by participants, predicted by agents, led to similarly improved BART performance in humans. Those who used a Bimodal strategy had significantly longer ITs of yellow balloons (*two-tailed t-test, t = 3.5 p = 0.011*; Fig. 3**e**), resulting in significantly higher scores (*two-tailed t test*, t = 9.1, *p < 0.001*; Fig. 3**f**). Interestingly, there was also no statistical difference in pop rate between the two behavioral classes of humans (*two-tailed t-test, t = −0.44, p = 0.66*; Fig. 3**g**). Together, these results demonstrate distinct classes of variably risk-adaptive behavior, predicted by randomly initialized deep RL agents, in human participants. One less successful strategy involves cautious exploration of the reward contingency space, whereas the more adaptive strategy involves underestimating gains for risky conditions while exploiting higher rewards in less risky conditions.

### Low-dimensional, factorized dynamical computations underlying risky choice in human neurons

To determine whether the structure of dynamical computation during human risky choices corresponded to the predictions from the deep RL agents, we recorded 597 well-isolated single units, going forward/henceforth referred to as neurons, from the medial frontal and temporal lobes of the participants while they carried out the BART (12.7 ± 7.8 units per session, Fig 4**a**). To measure the geometrical structure of dynamical computation that underlay each behavioral strategy in human neural ensembles, neurons were divided into two pseudoensembles corresponding to Bimodal (n = 473 units) and Explorer (n = 124 units) behavioral strategies. All neurons recorded from Bimodal participants were assigned to the Bimodal pseudoensemble and all neurons recorded from Explorer patients were assigned to the Explorer pseudoensemble. Initially, these neurons were analyzed to see if individual neurons encoded task-relevant variables. A Kruskal-Wallis test was done for each neuron by averaging the firing rates across a 1s *a priori* time window (0.2s – 1.2s). Then the χ^2^ value from the Kruskal-Wallis test was compared to a distribution of 10,000 permutations where firing rates were associated with shuffled trial labels. 4.2% of Bimodal neurons and 5.6% of Explorer neurons exhibited significant encoding of balloon size categories (Fig 4**b-e**). Therefore, at the level of firing rate encoding, we could not distinguish Bimodal from Explorer participants.

**Figure 4.**
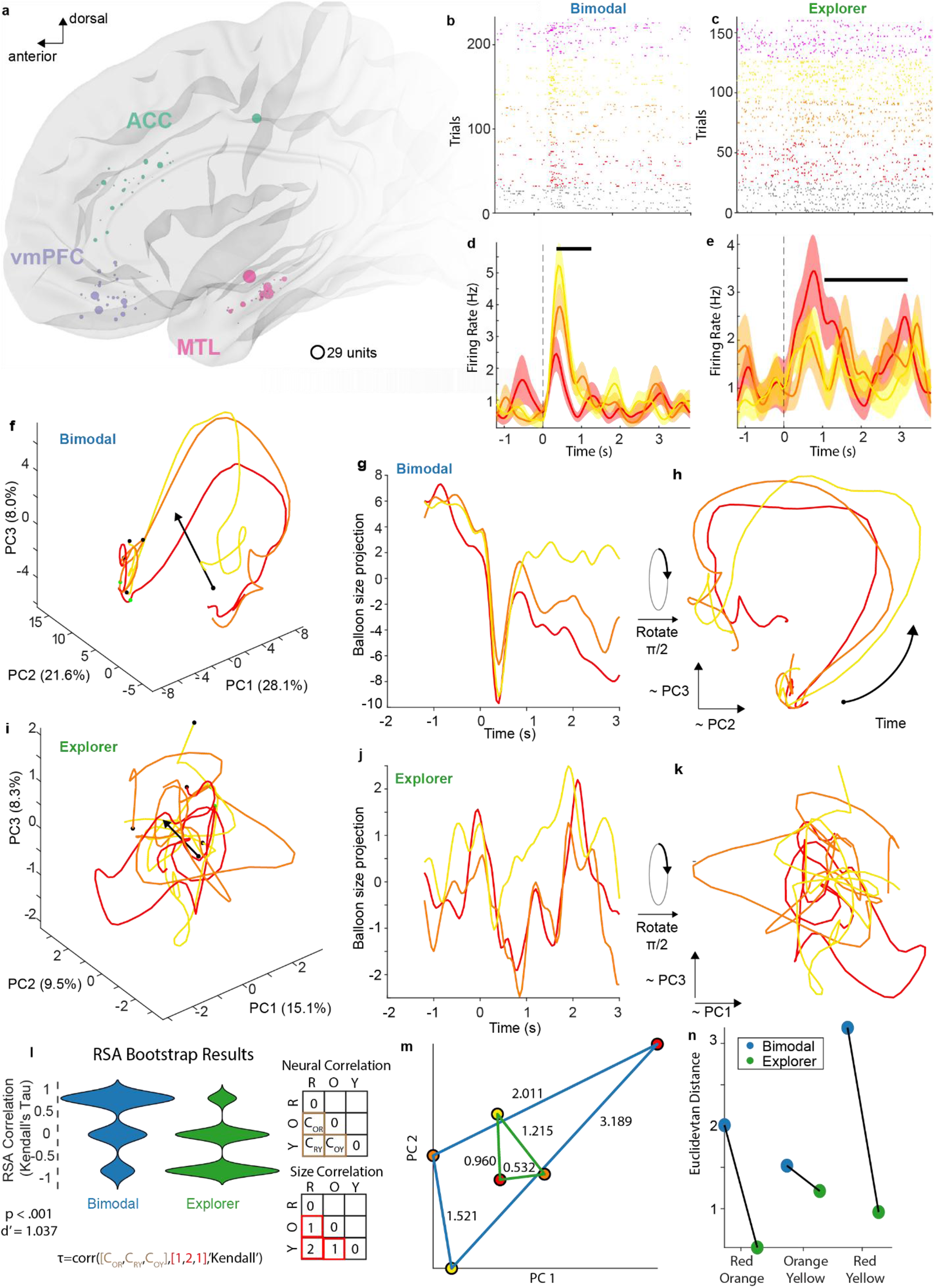
Human neuronal dynamical computation of risky choice. **a**, Single-unit (neuron) recording sites projected onto a common brain surface. Size of circle indicates how many units (neurons) were isolated from the microelectrode. The largest circle indicates 29 units were isolated, and fewer neurons were isolated from locations with smaller circles. (Green = ACC, Purple = MFC and Pink = MTL). **b-e,** Single-unit responses to BART. **b,d** Raster plot and accompanying PSTH for a unit (neuron) in the Bimodal pseudoensemble from vmPFC. **c,e** Raster plot and accompanying PSTH for a unit (neuron) in the Explorer pseudoensemble from MTL. (**d,e**) black line indicates period of significantly different firing rates (Wilcoxon Rank sum p < 0.05), **f-h,** Neural trajectories of the pseudoensembles of Bimodal units plotted in 3D PCA space, N_Bimodal Units_ = 473 (**f**), projected on to the balloon size axis (**g**), and rotated orthogonally relative to the balloon size axis (**h**). Black arrows show the decoded balloon size axes Black dots are the beginning of the recording window and green dot is when cue appears on screen. **i-k,** Same as **f-h**, but for Explorer units, N_Explorer Units_ = 124. **l,** (left) Violin plot of bootstrap (n = 10,000) RSA analysis. (right) Example calculation of a single Kendall’s Correlation, where the neural correlations of the different balloon sizes are correlated against the differences in actual balloon size. (Size matrix made by taking the absolute value of the difference between balloon sizes where Red = 1, Orange = 2 and Yellow = 3). **m,** Pairwise Euclidean distances between centroids of different trial types organized by behavioral category. **n,** Euclidean distances between centroids of PCA trajectories.

We next sought to test whether the structure of dynamical computation in each neuronal pseudoensemble reflected the distinct algorithms predicted by deep RL. To assess neural computation about participants’ expected outcomes, we examined firing rates aligned to the reward and risk predictive cue: the onset of the balloon. Firing rates were trial-averaged for each balloon color and organized into 3-D tensors (neurons x time x balloon color), which were unwrapped along the neuron dimension to create a data matrix for principal components analysis. We visualized the top three PCs of this matrix as neural trajectories for both Bimodal and Explorer pseudoensembles. As with the deep RL agents, we again estimated a predicted balloon size axis using partial least squares regression in 3-D PCA space.

We observed a remarkably similar structure in Bimodal human neural trajectories to those predicted by the recurrent layers of the deep RL agents, with trajectories leaving a pre-balloon fixed point to arc around an ordered balloon size axis that clearly segregated the predicted value of each balloon color (Fig. 4**f-h**). In contrast, neural trajectories from the Explorer group did not show such structure and were visually distinct from the deep RL Explorer agents (Fig. 4**i-k**). Upon projecting the PCA trajectories onto their respective balloon size axes, Bimodal pseudoensembles exhibited increasing coactivity for each balloon size (red, orange, and yellow) between the appearance of the balloon and the end of the cue-aligned window (Fig. 4**g**). A similar coactivity pattern was notably absent from the Explorer projections (Fig. 4**j**). In Bimodal participants, and predicted by Bimodal agents, the 2-D convex hulls of the dynamical arcs increased for balloons with larger mean inflations (red trajectory: 178.4, orange trajectory: 235.8, yellow trajectory: 250.2; Fig. 4**h**). More variance was explained by cue-aligned PCs in the Bimodal group than the Explorer group (PCs 1-3 explained 32.9% of the variance for Explorers and 57.7% for Bimodal.; Supp. Fig. 4A). This result demonstrates that humans using a Bimodal strategy exhibit a more robust predictive model of the task that, as in the agents, led to improved performance.

Similar findings were true in outcome-aligned pseudoensemble trajectories. For example, the first three Bimodal PCs explained more variance than the Explorer PCs (Supp. Fig. 5**e**). The 3-D Bimodal PCA trajectory projected along a size axis saw its coactivity increase until the outcome of the trial occurred, at which point it decreased precipitously (Supp. Fig. 5**d**). At outcome and when projected onto the size axis, the yellow balloons had the lowest coactivity level, while red had the highest, which is an inversion of the expected reward levels (Supp. Fig. 5**d**). Explorer pseudoensembles projected onto their size axis slightly increased their coactivity throughout the outcome-aligned analysis window, without any task-relevant structure (Supp. Fig. 5**c**). The cosine angle between the cue-aligned size axis and PC1 for Bimodal pseudoensembles was 114.0^⸰^ and 64.8^⸰^ for Explorers, which was analogous to the relationships observed in the deep RL agents.

Once it was established that the meaningful visual differences in neural trajectories were not the result of the group size disparities, we undertook Representational Similarity Analysis (RSA) to quantify key features of the pseudoensemble neural representations at the level of averaged firing rates, rather than dynamical representations. RSA revealed that the Bimodal activity patterns significantly correlated with predicted balloon sizes (*bootstrap τ = 0.32*), whereas Explorer patterns showed negative correlation with predicted balloon sizes (*τ = −0.28; Wilcoxon-Rank sum test, p < 0.001, Cohen’s d = 1.03*; Fig. 4**l**). This indicates that Bimodal neurons encode balloon size in a manner consistent with physical size relationships, while Explorer neurons exhibit inverted or absent size representations. Pairwise centroid distances revealed divergent geometries: Bimodal participants expanded Red-Orange separation (2.3× Explorer distance) while compressing Orange-Yellow distances (0.4× Explorer; Fig. 4**n**). This geometry suggests a binary ‘avoid vs. pursue’ decision boundary: as in the deep RL agents, as a group, humans using a Bimodal strategy encoded Red as categorically distinct from Orange and Yellow (Stop Red, Inflate Orange/Yellow), whereas those using an Explorer strategy collapse Red and Orange representations, while separating Yellow (Stop Red/Orange, Inflate Yellow). Thus, Bimodal humans compress their state space around behaviorally relevant boundaries, treating medium and high-value balloons as a unified “inflate-worthy” category, resulting in the risk-adaptive Bimodal human behavior paralleling the dynamics and behavior found in Bimodal agents.

While we initially analyzed all neurons together, we noticed brain area-specific differences in balloon-aligned coactivity. Upon examining trial-averaged firing rates of the two pseudoensembles by brain area, the anterior cingulate cortex (ACC) showed a clear reward prediction signal in the Bimodal pseudoensemble (Supp. Fig. 4**d**). Interestingly, when the sum of squared loading was calculated for each brain area only about 18.5% of the variance explained in PC1 in the Bimodal group came from ACC; for Explorers, the ACC variance explained was 56.0%. The brain area that accounted for the largest amount of variance in PC1 for the Bimodal group was vmPFC (72.9%). For the Bimodal group, ACC accounted for the largest amount of variance in PC2 (63.9%; Supp. Figs. 4**n-o**). To exclude the possibility that the disparity in geometries between behavioral classes was the result of the differences in number of neurons between the two subpopulations or the differences in proportion of neurons in different brain areas, two control analyses were conducted. First, the Procrustes distance between a random, balanced subsample (without replacement) of Bimodal units and the original Explorer geometry was calculated 10,000 times. The true Procrustes distance between the geometries of Bimodal and Explorer pseudoensembles was determined to be significantly smaller than expected by chance (*permutation test, p = 0.01*). Next, the proportion of neurons in each of the four brain areas from which the units were recorded were held constant, such that the subsample of the Bimodal subpopulation had the same number of units as the original Explorer population and had the same number of units from each of the brain areas as the original Explorer subpopulation. Again, the true Procrustes distance was smaller than expected by chance (*permutation test, p = 0.009*). These controls indicated that differences between Bimodal and Explorer geometries could neither be explained by differences in overall sample sizes, nor sample sizes from each brain area and resolved concerns that a higher proportion of ACC units did not lead to more organized geometries (Supp. Fig. 4**i-j**).

## Discussion

In this study, we leveraged deep RL to predict distinct classes of algorithms and their underlying dynamical computations in human neuronal ensembles that subserve behavior during a complex, ethologically valid risky decision-making task.^30^ We found that deep RL agents spontaneously differentiated into distinct behavioral phenotypes based on the proportion of unit activity types in their recurrent layers, which we called Bimodal and Explorer. Human participants performing the BART exhibited analogous behavioral strategies, with the Bimodal group achieving significantly higher performance in both artificial and biological agents. Our central finding is that this behavioral convergence is supported by a convergence in neural geometry: the neuronal ensemble dynamics underlying risk-adaptive human behavior shared the same untangled low-dimensional trajectories arcing around a balloon size axis predicting expected reward, as discovered in Bimodal agents. These results suggest that specific neural population geometries may be critical for supporting learned inference required for adaptive behavior in conditions with uncertain outcomes.

Notably, the predictions from the deep RL algorithms arose from marked behavioral diversity, stemming largely from random initializations and different task parametrizations. There is accumulating evidence that deep RL techniques can generate useful predictions for individual behavior and its underlying neural substrates.^31,32^ This framework has thus proved beneficial for generating predictions about the diversity of neural algorithms, and their underlying geometries, resulting from varied initial conditions, and training parameters.^33^

Through analogous analysis of artificial and biological neural networks, we were able to test predictions about the neural algorithms employed to carry out the BART, and their associated underlying dynamical computations. Procrustes distance, a measure of how similar two multidimensional geometries are to one another, confirmed that the geometries of the human Bimodal trajectories were not simply due to the differences in sample size but likely reflect a more robust internal model in the brains of Bimodal participants. This model is characterized by untangled, or factorized, representations allowing for organized readout of neural activity for implementing behavior. These results further support growing evidence in the “Neuro-AI” field that artificial networks can serve as models to unpack the dynamical computations underlying high-level cognitive processes, not just sensory representations.^15,23,34–36^ Specifically, these results show how the human brain compresses complex task state spaces (like the probabilistic growth of a balloon tied to rewards) into low-dimensional dynamics on manifolds that simplify the readout for downstream motor or decision circuits. Additionally, these brain areas were originally targeted for their respective contributions to cognitive control. On average, the size axis derived from the Bimodal pseudoensemble was (anti) parallel with PC1. We interpreted that the larger vmPFC contribution to PC 1 in the Bimodal pseudoensemble indicated that participants using a Bimodal strategy were potentially prospectively considering each balloon’s potential value or risk categories as they inflated each balloon. Alternatively, the increased variance contribution from ACC in the Explorer group might suggest they were monitoring for reward prediction errors more than the Bimodal group. These qualitative differences in brain area contributions suggest that Bimodal humans may be creating a model of the BART task and are optimizing their behavior based on their understanding of the underlying task structure. Conversely, Explorers may be motivated by recent reward or outcome patterns. This dichotomy mirrors the well-established model-based and model-free inference described in the RL and cognitive science literature^37–41^.

There are several limitations that apply to this study. First, our human cohort consisted of drug-resistant epilepsy patients, and while our sample size of 44 patients is large for this field, the electrode coverage was sparse and limited to specific regions (ACC, vmPFC, MTL), as determined by clinical necessity. The yield of units is smaller than in some other papers, but we prioritized having patients only run one session of BART and only two to three BF electrodes were implanted in each patient. While we verified that anatomical sampling did not drive the geometric differences, we cannot rule out the contributions unsampled brain regions. Second, the PPO agents are abstractions that do not simulate biological plasticity or neuromodulation. Future work could incorporate more biologically plausible learning rules or spiking neural networks to see if they converge on similar choice-predictive manifolds. Despite these limitations, our results provide a compelling demonstration that aligning the latent spaces of biological and artificial agents can reveal the shared algorithmic and computational principles of decision-making.

## Materials & Methods

### Meta-BART for deep RL agents

Meta-BART is a variation of the human BART, where digital agents were presented “balloons” for a number of episodes. Each episode consisted of 50 balloons. At the start of each episode, a mean balloon size, *µ*, was drawn from a uniform distribution, U(0.2, 1), and the maximum size of the individual balloons in the episode was drawn from N(µ,0.05). Agents were trained on multiple variants of the meta-BART environment. Variants are distinguished by three variables. First, agents may or may not have been given explicit reward information as part of their observation. Second, reward was calculated as *r = s^h^*, where *s* is the size of the balloon when banked and *h* ∈ (1, 1.2, 1.5, 1.7 2) is a training variable that shapes rewards to more strongly reward larger balloons than smaller balloons as *h* increases, encouraging the development of risk-seeking agents. Finally, when a balloon popped, a punishment (0, −0.1, −0.2, −0.4) could be given. Agents were trained in each possible combination of these variables, leading to 40 training conditions, (Supp. Fig. 1**b**).

For each condition, 10 agents of different random seeds were trained, leading to a total of 400 agents were developed for analysis in this work. At each time step, the agent had two actions available: start/stop or wait. Once inflation began, the balloon inflated at a fixed rate of 0.05 units per time step until the agent stopped inflation or the balloon size reached its maximum and popped. If the agent used the stop action, it received a reward related to the size of the balloon at stopping. If the balloon popped, a punishment was applied according to the punishment parameter.

### Training deep RL agents

Training deep RL agents followed a standard proximal policy optimization (PPO) method.^42^ Each agent’s neural network consisted of a shared feed-forward layer, a RNN-Layer, which connected to actor and critic feed-forward layers to output the policy (π) and value estimations (V), respectively. Every layer had N = 64 nodes and the weights of each agent network were randomly initialized. The agent neural network nodes all used hyperbolic tangent activation functions. The agents observations, o_t_ ∈ ℝ^3^, were vectors formed by the size of the balloon, the previous action a_t-1_ (1 if the button was pressed, 0 otherwise) and the previous reward r_t-1_.

To collect training data, the policy (π) was run in 16 parallel copies of the simulated meta-BART task.^43^ Parallel environments were allowed to each have different (µ) meta conditions, as is standard in this training paradigm. State, action, reward tuples (s_t_, a_t_, r_t_, s_t+1_) were collected from each simulation for 256 steps, cumulating to a total batch size of 4096 steps. PPO was used to optimize the neural network on the batch of data, and then the collection and training loop repeated. Agent networks were frozen and saved every 30 batches of data (roughly 120,000 time steps), which we refer to henceforth as *checkpoints*, up to a maximum of 240 batches or 8 checkpoints (roughly 1 million time steps).

### Agent evaluation

Agent performance was evaluated across an evaluation suite of fixed conditions µ ∈ (0.2, 0.25, …., 0.95, 1), for a total of 17 meta conditions. As in training, 50 balloons were drawn for each meta condition that agents were tasked with inflating and stopping, to earn points. The evaluation runs were seeded such that while balloon sizes are drawn from N(µ,0.05) the actual sizes were consistent for every run. However, actions were drawn from policy outputs stochastically, which is contrary to typical evaluations in RL where the highest probability action is deterministically taken at every time step. Only one saved checkpoint per agent was used to analyze their behaviors and internal representations. Starting from checkpoint 1, agents were tested on the evaluation suite at every checkpoint and their total banked balloon size was recorded. The first checkpoint that exceeded a total banked size of 275 on the 17 episodes for each agent was saved for further analysis. Otherwise, if an agent never reached the 275 threshold across the tested checkpoints, the best performing agent was used.

Symmetry induced by the hyperbolic tangent function makes it so that the same information can be propagated from a node to the next layer whether the node uses negative or positive activities by changing the sign of the forward weight. This is an important consideration when using k-means clustering on node activities to find similarly activating nodes. We were more interested in the shape of node activities in terms of the information they can encode than their absolute values. To compare agent node dynamics, we went through a multi-step normalization process on the collection of node activities across all episodes for each agent, *{(z_i,t_)}_i= 0,…,N_*, before applying k-means on these node activities. First, all activities were normalized to have zero mean and unit variance 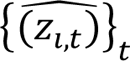. Next, every activation was duplicated and made negative, so the collection becomes 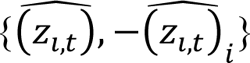. K-means was applied to this collection selecting 2*k* clusters where *k* was the desired final number of clusters. This process resulted in paired clusters that reflected each other along the x-axis, due to duplicated activations. K = 12 was selected based on visually inspecting the elbow. We called two clusters a *pair* if when one is multiplied by −1, they had a resulting correlation coefficient greater than 0.9 and arbitrarily selected one cluster from the pair to keep, resulting in cluster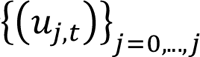. Increasing the number of clusters did not appear to generate cluster centers with new features in the dynamics.

Finally, with these reduced clusters, redundant node activities were removed that were added in the duplication step. To do this, for each node we found the minimal distance between either positive or negative node activity, 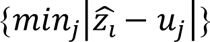 and 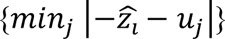. The minimum distance gave us the cluster, *j*, that the node is assigned to, as well as the orientation of the node activity (depending on whether the positive or negative activity was used), which could be used if one were interested in comparing node time course shapes to each other in a consistent way. Only the cluster centers and node cluster labels were used in this work. This process resulted in six (k = 6) remaining clusters that we ultimately used to classify agent identities.

### Human participants and ethics statement

Participants were adult drug-resistant epilepsy patients (n = 44 (23 female); M = 35.3 ± 12.2 years of age) undergoing intracranial monitoring as part of neurosurgical treatment for epilepsy. These patients were implanted with stereo-electroencephalographic (sEEG) depth electrodes, a subset of which contained microwires extending from their tips.^44^ All participants provided informed consent prior to microelectrode implantation and The University of Utah Institutional Review Board approved this study.

### Human behavior on the Balloon Analog Risk Task (BART)

BART is a validated cognitive-behavioral paradigm that can measure both risk-taking behavior and choice impulsivity.^24^ Each task session began with researchers instructing patients on the general task structure with the explicit goal to collect as many points as possible during the session. The relationship between balloon color and potential reward were explained, as well as the passive trial indicator. After receiving the instructions, participants completed more than 200 trials of BART while the electrical activity from intracranial microwires was recorded.

During each BART trial, participants press a button on a video game controller to begin inflating a computerized balloon that they observed on a computer monitor that was placed in front of them. Participants pressed the same button on a game controller to both start and stop the balloon’s inflation. If the inflation was stopped prior to the balloon reaching its maximum size, patients received points linearly related to the size of the balloon. If the balloon instead popped, the participants neither received nor lost any points. BART contained both active and passive trials. Passive trials included both rewarded or unrewarded balloons while participants always had an opportunity to earn a reward on active trials. Balloons appeared on the screen in one of four colors: gray, red, orange and yellow. All the uninflated balloons appeared as the same size and their maximum size was drawn from respective normal distributions and standard deviations. Gray balloons were never rewarded. Red balloons had a smaller maximum size than orange balloons, and orange balloons smaller than yellow balloons. Thus, there were five reward categories of balloons: gray (unrewarded passive trials), yellow, orange, red (rewarded active trials) and what are represented as pink (rewarded passive trials with yellow, orange, and red colors). The color of balloon, and the presence or absence of an indicator of a passive trial cued patients to the potential for obtaining reward on each trial.

BART was implemented in PsychToolbox for Matlab.^45,46^ Participants registered their behavioral responses via a USB game controller (Logitech G F310). Task events and patient behavior were synchronized via digital codes sent from the task control software to a PCI-express card in the task control computer with sub-millisecond precision.

### Electrophysiology

Single-unit activity was recorded from patients using Behnke-Fried (Adtech Medical; Racine WI) microwires extending from the distal tip of two to three of the patients’ clinical macroelectrodes on a Blackrock data acquisition system (Blackrock Neurotech; Salt Lake City, UT) with a sampling rate of 30 kHz.^44^ Single units were isolated by bandpass filtering the signal from the microwires between 0.25 and 7.5 kHz and sorting waveforms that crossed −3.5 times the mean root squared of the filtered signal using a semi-supervised process based on t-distributed expectation maximization on the waveforms’ first three principal components via Offline Sorter spike sorting software (Plexon, Inc.; Dallas, TX).^47^

Prior work has shown that midbrain dopaminergic firing rates peak between 200-600 ms after the reward cue.^8,11^ The windows of this analysis were extended due to recording locations being downstream from these midbrain areas. To allow comparisons between different trial types a cue-aligned window began 1.2 s prior to the balloon appearing on screen and ended 3 s after. Our hypothesis was that the most reward-relevant signals would occur 200 – 1,200 ms after the reward predictive cue, or the outcome. A Kruskal-Wallis test was done for each cell to determine if it was encoding trial type by firing rate. Firing rates were averaged across a 1s *a priori* time window (0.2s – 1.2s). Then the χ^2^ value from the Kruskal-Wallis test was compared to a distribution of 10,000 permutations where firing rates were compared to shuffled labels.

### Electrode localization

Electrode locations were determined from co-registered pre-operative magnetic resonance imaging (MRI) and postoperative computed tomography (CT) using the LeGUI software package.^48^ Units were grouped into one of three gross anatomical regions based on where their microwires terminated: anterior cingulate cortex (ACC), ventromedial prefrontal cortex (vmPFC) and mesial temporal lobe (MTL). See Figure 4 for recording locations.

### Classification of behavioral strategies

We wanted to compare human behavioral strategies to those identified by the agents. Initially, only behavioral measures were used to classify the human participants. As in the artificial agents, participants were identified as ‘Bimodal’ if the difference between their mean inflation time (IT) for red balloons and their mean IT for orange balloons was three times greater or smaller than the difference between their mean IT for orange balloons and their mean IT for yellow balloons. ‘Explorer’ labels were assigned to any participant whose smallest balloon was within 80% of their first banked balloon and their largest balloon was within 120% of their final banked balloon. Due to the different maximum possible sizes for each color of the balloon, the trajectory of balloon sizes was examined for each color independently. Finally, participants who fit into neither category were classified as ‘Neither’. Since the definitions of ‘Bimodal’ and ‘Explorer’ were not mutually exclusive, we used k-means (k=3) on a higher-dimensional behavioral matrix including their total score, pop rate, and mean inflation time on active trials for each of the balloon colors. The behavioral classifications from k-means were used throughout the analysis.

### Comparing geometries

Pseudopopulations of units were built such that units from participants in the ‘Bimodal’ group went into the ‘Bimodal’ pseudopopulation, while units from the ‘Explorer’ participants went into the ‘Explorer’ pseudopopulation. Two dimensionality reduction methods were used to determine neuronal population geometry. Concatenated, trial-averaged Principal Component Analysis (PCA) was done on these pseudopopulations.^19^ PCA was used to compare both the agentic nodal representations as well as the different neuronal representations of the different behavioral factions within the patient population. Agentic trials consisting of only two time points were not included in the visualization and subsequent analysis because these trials consisted of the agent beginning the balloon inflation at the first time step, and then immediately ceasing inflation at the second time step. For each selected balloon mean, the minimum number of time points in each trial was included. Node activity exceeding this minimum number of time points was discarded. The built-in Matlab function ‘procrustes,’ using the default Euclidian Distance, was used to compute the differences between the ‘Explorer’ and ‘Bimodal’ geometries using 5 PCs.

To determine a size axis for each of the agents a partial least squares (PLS) regression was done using scikit-learn’s cross decomposition toolbox.^49^ Chosen mean balloon size (µ) was regressed onto the activations of the recurrent layer. The size axis was defined as the x_weights of the PLS regression. This initial size axis was multiplied by the trial averaged PCA components to orient the size axis into PC space, and to determine its angle difference from PC axes. To help orient the subsequent analysis, the size axis was given a direction by making sure whichever side of the axis the µ = 1 condition was positive.

For the size axis of humans, a similar process was followed with the only modification being that the average IT for each balloon condition was regressed onto the respective firing rates. One additional difference is that each agent was fit with their own size axis while only one size axis was generated for each pseudopopulation of human participants.

The sum of squared loadings for each unit in their respective brain areas for each PC was calculated to quantify each area’s contribution.

To compare the distributions of neural trajectories along the balloon size axes, PCA trajectories were projected onto the size axis for each agent. This led trajectories to appear as relatively straight lines evolving in time, distributed along the one-dimensional balloon size axis. Mean values for each trajectory, corresponding to each *µ*, were then z-scored for each agent respectively. Once z-scored, these values were visualized with histograms and a Hartigan and Hartigan Dip test was used to determine if and which resulting distributions exhibited a Bimodal distribution.

Cosine angles for the agents were calculated by reflecting all angles greater than 90^⸰^ (π/2) so that they were between 0 and 90^⸰^ and then calculated the means and standard deviations. Without the reflection the means were very similar (Bimodal = 98.5^⸰^ ± 66.7^⸰^; Explorer = 88.7^⸰^ ± 36.5^⸰^.

### Quantification and Statistical Analysis

All analysis was done using custom scripts in Matlab (Mathworks; Natick MA) and Python (pyOrg).^49,50^ Criterion was set at 0.05. All code and data are publicly available.

Non-parametric permutation tests (n = 10,000) were used to test the significance of the Procrustes distance between the two geometries. Two different simulations were run; the first test was done by randomly sampling 124 Bimodal units and comparing their geometry to that of the actual Explorer geometry using Procrustes Distance. The second test was done by sampling 124 Bimodal units but maintaining the proportion of units from each of the four brain areas and comparing it to the actual Explorer geometry.

To quantify the geometric structure of neural population activity, we computed three metrics: Representational Similarity Analysis (RSA), orthogonality, and separability. For RSA, we constructed neural representational dissimilarity matrices (RDMs) by computing Pearson correlation distances between population activity patterns for each balloon condition (Red, Orange, Yellow) averaged across the cue-aligned window (0.2-1.2 s post-cue). Neural RDMs were correlated with a model RDM encoding physical balloon size relationships using Kendall’s tau. To control for unequal unit counts between groups (Bimodal: 473 units, Explorer: 124 units), both groups were bootstrapped to 124 units with replacement for 10,000 iterations, generating distributions for statistical comparison.

Pairwise Euclidean distances between condition centroids were calculated in PC space to visualize representational warping. Centroids were centered to the origin to remove overall position differences and facilitate geometric comparison. Full pseudoensemble trajectory values are reported for reference, but statistical comparisons used bootstrapped subsamples to control for unit count disparities.

## Data & Code Availability Statements

Supplementary Information is available for this paper.

Code available <u>here</u>

Data available <u>here</u>

## Author Contributions

T.A.P. collected and analyzed human data as well as agent data. A.L. designed the artificial agents, and the metaBART environment as well as analyzed agent data. R.L.C., N.S. and T.S.D collected human data and aided in analysis. B.K., J.D.R, S.R. and B.S. implanted electrodes into neurosurgical patients, enabling the collection of human data. A.B and E.H.S designed the study, discussed the findings. T.A.P., A.L, A.B., E.H.S wrote the paper. All authors reviewed and edited the paper.

## Acknowledgments

We thank the patients and their families for taking the time to participate in research. This work was supported by a grant from the National Institutes of Mental Health (R01MH128187).

---

**Supplementary Figure 1.**
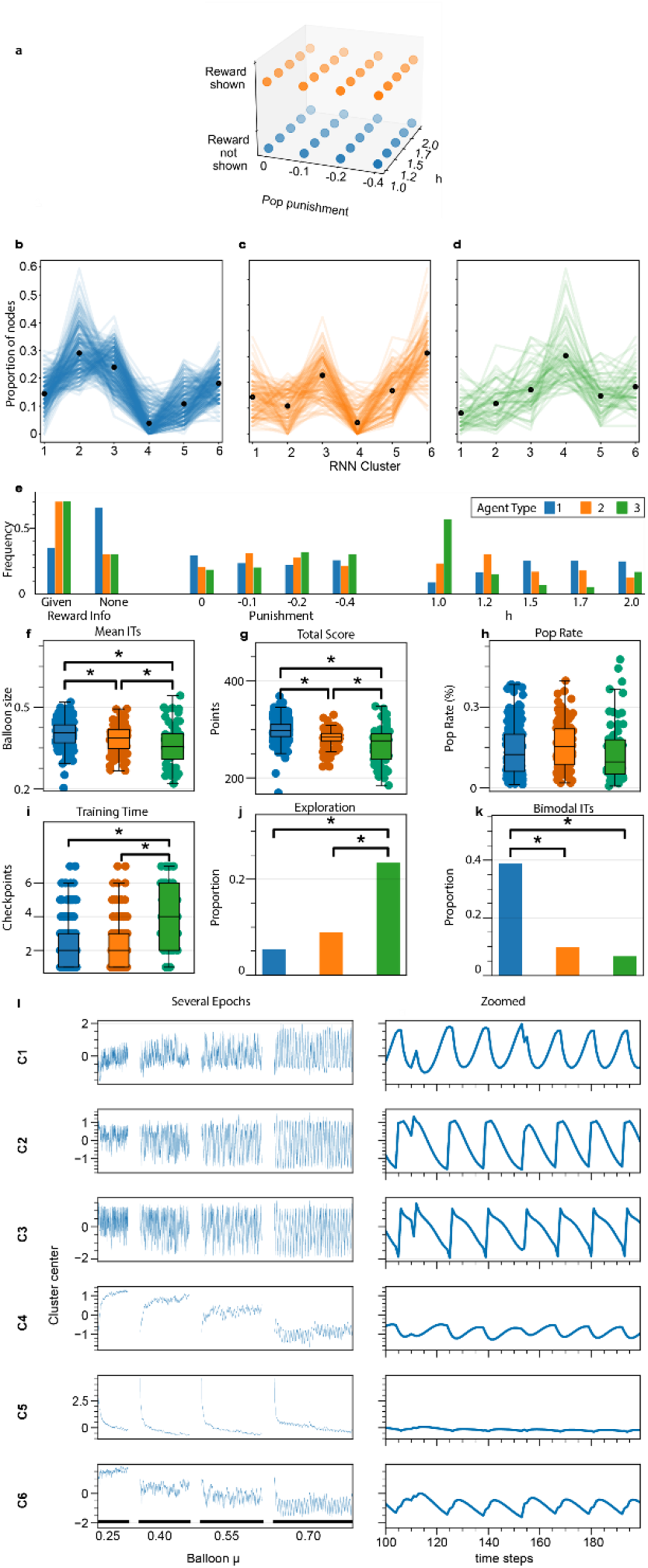
Representative activations for deep RL agent node clusters. **a**, training parameters for the agents. **b-d,** Proportion of nodes from each of the three agent types. Type 1 (**b**) are very similar to Type 2 (**c**), with the exception of relative proportion of node type 2. Type 3 agents are pictured in (**d**). **b,d**, similar to Fig 1. **e,** Frequency of agent types that developed based on different task parameters. Type 1 = Bimodal, Type 3 = Explorer. **f**, Mean inflation time for each of the agents, paired t-tests. **g**, Total score for each agent type, paired t-tests. **h**, Pop rate for each agent type, **i**, Number of training checkpoints until performance level achieved, paired t-tests. **j**, Exploration metric for each type of agent type, paired t-tests **k,** Proportion of agents with bimodal inflation times, paired t-tests. **l,** Examples of six clusters of node activities. The left column shows node activity across several epochs. The right column shows node activity across approximately 100 time steps. *, **, *** indicate statistical significance (p<0.05, p<0.01, p<0.001)

**Supplemental Figure 2.**
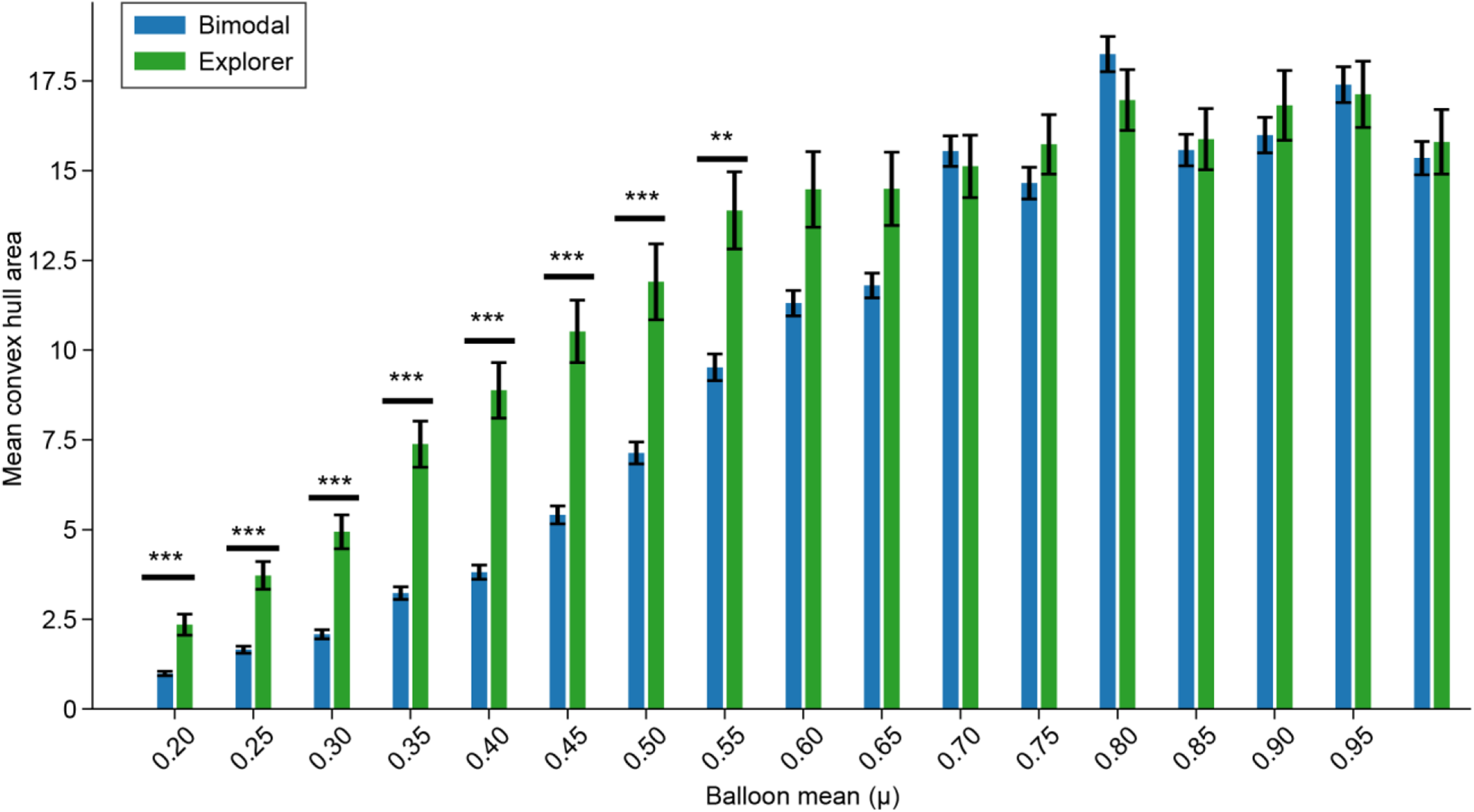
Surface area of neural trajectory arcs in deep RL agents. Measures of the area of the convex hull of the trajectories in PCA space of Bimodal and Explorer agents. Error bars are SEM. *, **, *** indicate statistical significance (p<0.05, p<0.01, p<0.001).

**Supplementary Figure 3.**
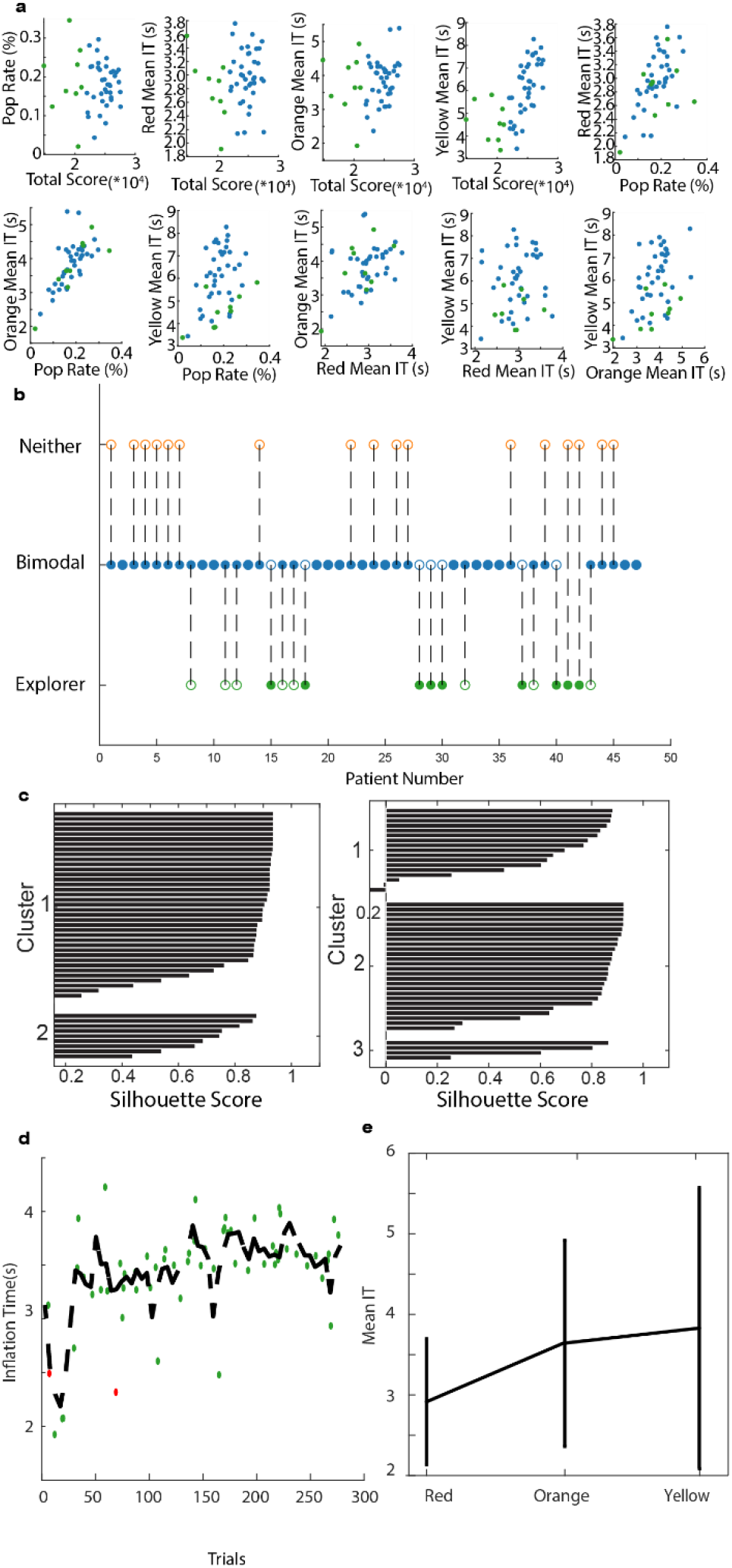
Clustering human risky choice behavior. **a**, Behavioral measures on which k-means clustering was done. blue corresponds to Bimodal, green corresponds to Explorer **b,** Open circles indicate the original assignment of the patient based solely on behavioral metrics. Closed circles indicate the final categorization of the patient based on k-means clustering. **c,** Silhouette scores of k-means clustering. k = 2 led to better silhouette scores than k = 3, so k = 2 was used and the ‘Neither’ categorization was dropped. **d**, an example Bimodal patient where yellow balloon sizes were learned after only a few trials. **e**, Mean inflation times for red, orange and yellow balloons respectively for an example Explorer human.

**Supplemental Figure 4.**
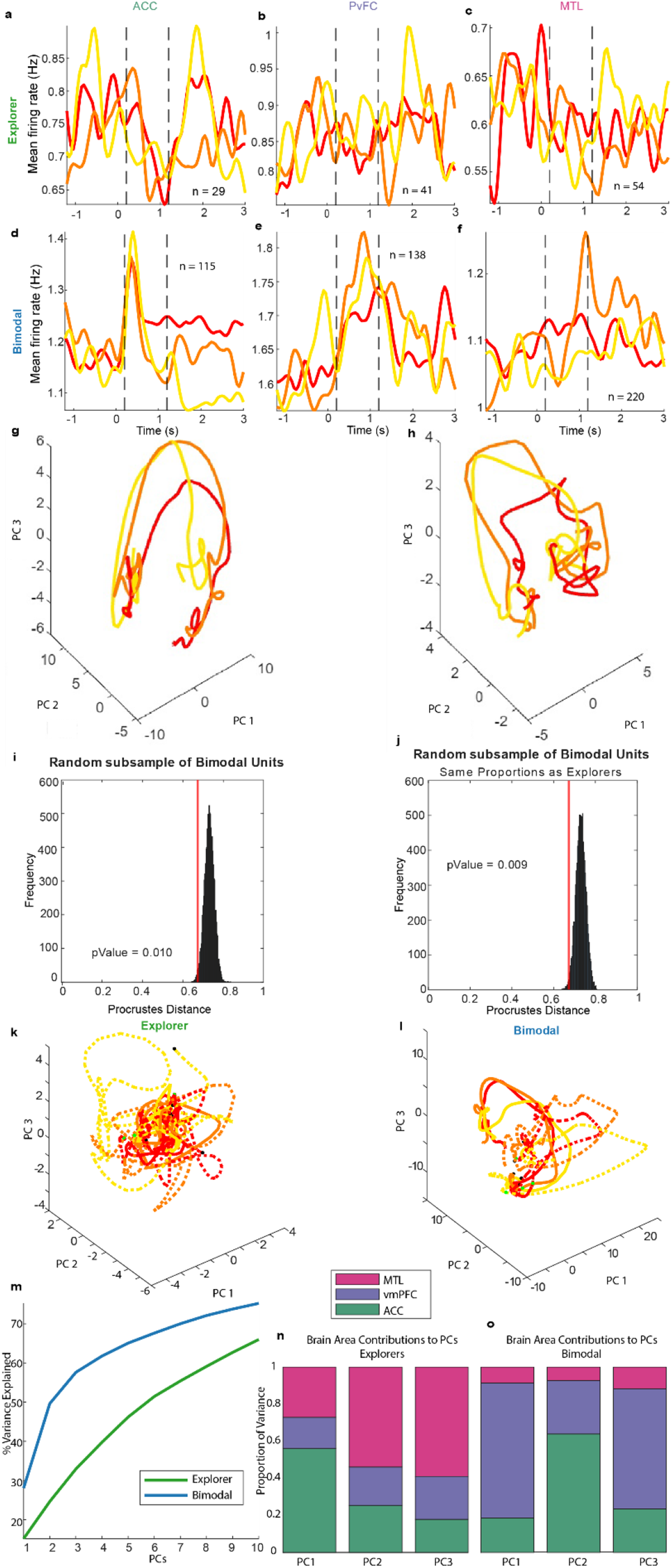
Features of human neuron dynamical computation. **a-f**, Trial-averaged firing rates by behavioral class and brain region. Dashed lines mark the 200 ms −1200 ms *a priori* window. **g-h,** PCA trajectories of neural trajectories from subsampled Bimodal pseudopopulation. The relative proportions of units sampled from each brain area were held consistent with the Explorer pseudopopulation. **i,j,** Permutation tests for Procrustes Distance metric comparing the geometries of subsampled neurons from the Bimodal and Explorer groups. **k,** the Bimodal population of units was subsampled without replacement such that the subpopulation had the same number of units as the Explorer group. **l,** the Bimodal population of units was subsampled such that the subpopulation had the same number of units as the Explorer group and that the proportion of units from each brain area matched (i.e. the subpopulation of Bimodal units contained exactly 29 units). **k-l,** 3D cue-aligned PCA trajectories for the pseudopopulations of Explorer (**k**) and Bimodal (**l**) with active trials represented as solid lines and passive trials represented as dashed lines. **m**, Variance explained for PCA on cue-aligned firing rates for Bimodal and Explorer humans. **n-o**, Proportion of variance explained by each brain area grouped by PC for Explorers (**n**) and Bimodal (**o**) pseudoensembles (sum of squared loadings).

**Supplemental Figure 5.**
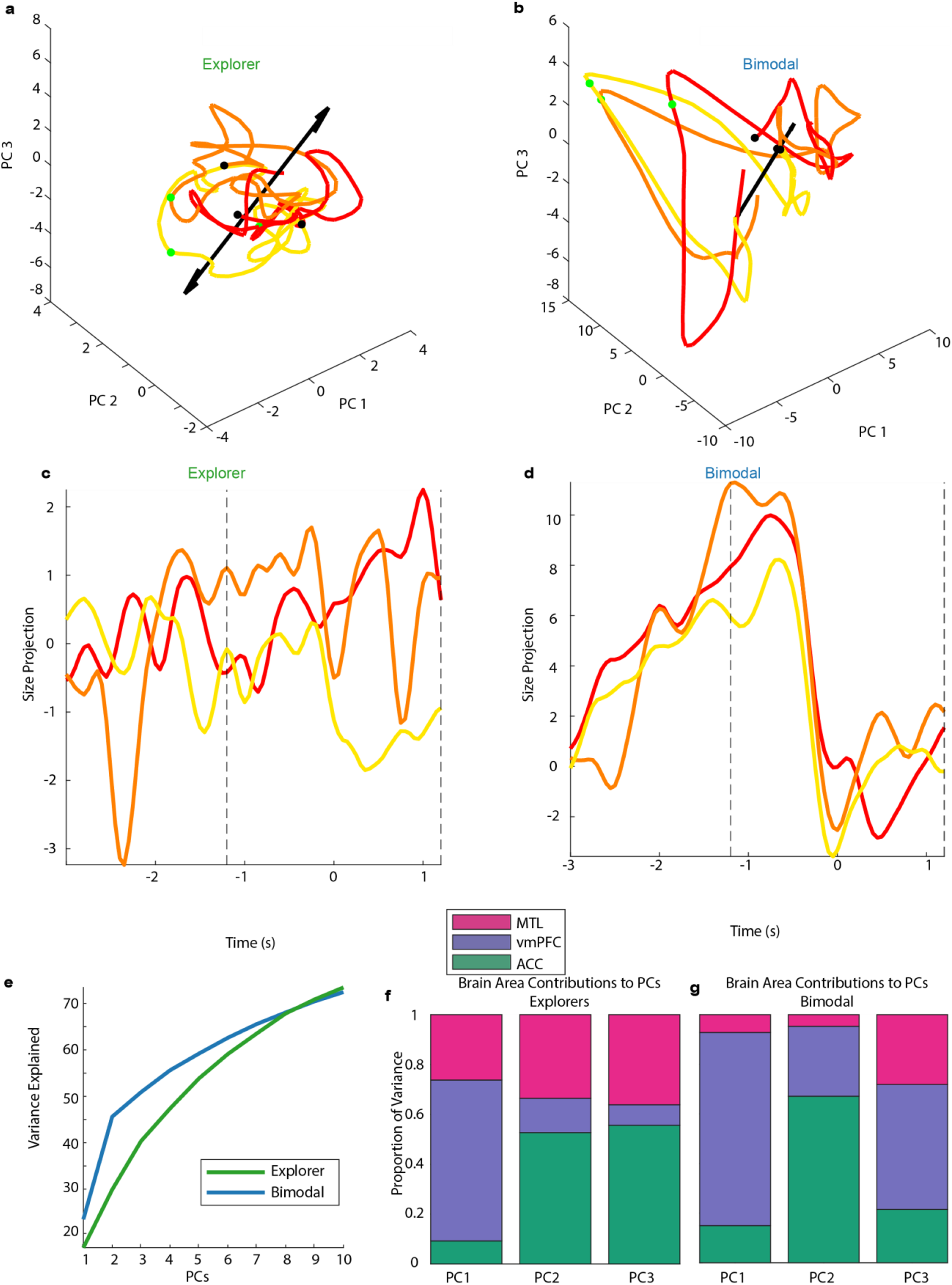
Outcome-aligned neural trajectories. **a,b** Outcome aligned neural trajectories in 3D PCA space for humans using Explorer (**a**) and Bimodal (**b**) strategies. Black arrows are decoded size axes, black dots are beginning of recording window and green dot is when outcome occurs. **c-d,** Trajectories projected onto the Size Axis. Dotted lines bracket (± 1200 ms)the moment that outcome occurs. **e,** Variance explained for 10 PCs. **f-g,** Proportion of variance explained by each brain area grouped by PC for Explorers (**f**) and Bimodal (**g**) pseudoensembles (sum of squared loadings).

## References

1. Behrens, T. E. J., Woolrich, M. W., Walton, M. E. & Rushworth, M. F. S. Learning the value of information in an uncertain world. Nat. Neurosci. 10, 1214–1221 (2007).

2. Rosenbaum, G. M. & Hartley, C. A. Developmental perspectives on risky and impulsive choice. Philos. Trans. R. Soc. B Biol. Sci. 374, 20180133 (2019).

3. Read, R. W. et al. A study of impulsivity and adverse childhood experiences in a population health setting. Front. Public Health 12, (2024).

4. Everitt, B. J. & Robbins, T. W. Neural systems of reinforcement for drug addiction: from actions to habits to compulsion. Nat. Neurosci. 8, 1481–1489 (2005).

5. Sutton, R. S. & Barto, A. G. Reinforcement Learning: An Introduction. (The MIT Press, Cambridge, Massachusetts, 2018).

6. Rummery, G. & Niranjan, M. On-Line Q-Learning Using Connectionist Systems. Tech. Rep. CUEDF-INFENGTR 166 (1994).

7. Bellemare, M. G., Dabney, W. & Munos, R. A distributional perspective on reinforcement learning. in Proceedings of the 34th International Conference on Machine Learning-Volume 70 449–458 (JMLR.org, 2017).

8. Dabney, W. et al. A distributional code for value in dopamine-based reinforcement learning. Nature 577, 671–675 (2020).

9. Schütt, H. H., Kim, D. & Ma, W. J. Reward prediction error neurons implement an efficient code for reward. Nat. Neurosci. 27, 1333–1339 (2024).

10. Sousa, M. et al. A multidimensional distributional map of future reward in dopamine neurons. Nature 642, 691–699 (2025).

11. Muller, T. H. et al. Distributional reinforcement learning in prefrontal cortex. Nat. Neurosci. (2024) doi:10.1038/s41593-023-01535-w.

12. Jurewicz, K., Sleezer, B. J., Mehta, P. S., Hayden, B. Y. & Ebitz, R. B. Irrational choices via a curvilinear representational geometry for value. 2022.03.31.486635 Preprint at 10.1101/2022.03.31.486635 (2022).

13. Watts, T. W., Duncan, G. J. & Quan, H. Revisiting the Marshmallow Test: A Conceptual Replication Investigating Links Between Early Delay of Gratification and Later Outcomes. Psychol. Sci. 29, 1159–1177 (2018).

14. Chung, S. & Abbott, L. F. Neural population geometry: An approach for understanding biological and artificial neural networks. Curr. Opin. Neurobiol. 70, 137–144 (2021).

15. Gallego, J. A., Perich, M. G., Miller, L. E. & Solla, S. A. Neural Manifolds for the Control of Movement. Neuron 94, 978–984 (2017).

16. Russo, A. A. et al. Motor Cortex Embeds Muscle-like Commands in an Untangled Population Response. Neuron 97, 953–966.e8 (2018).

17. Fu, Z. et al. The geometry of domain-general performance monitoring in the human medial frontal cortex. Science 376, eabm9922 (2022).

18. Feather, J. & Chung, S. Unveiling the benefits of multitasking in disentangled representation formation. Trends Cogn. Sci. 27, 699–701 (2023).

19. Vyas, S., Golub, M. D., Sussillo, D. & Shenoy, K. V. Computation Through Neural Population Dynamics. Annu. Rev. Neurosci. 43, 249–275 (2020).

20. Wang, J. Prefrontal Cortex as a Meta-Reinforcement Learning System. Reinf. Learn.

21. Hassabis, D., Kumaran, D., Summerfield, C. & Botvinick, M. Neuroscience-Inspired Artificial Intelligence. Neuron 95, 245–258 (2017).

22. Sussillo, D., Churchland, M. M., Kaufman, M. T. & Shenoy, K. V. A neural network that finds a naturalistic solution for the production of muscle activity. Nat. Neurosci. 18, 1025–1033 (2015).

23. Sohn, H., Narain, D., Meirhaeghe, N. & Jazayeri, M. Bayesian Computation through Cortical Latent Dynamics. Neuron 103, 934–947.e5 (2019).

24. Lejuez, C. W. et al. Evaluation of a behavioral measure of risk taking: The Balloon Analogue Risk Task (BART). J. Exp. Psychol. Appl. 8, 75–84 (2002).

25. Hunt, M. K., Hopko, D. R., Bare, R., Lejuez, C. W. & Robinson, E. V. Construct Validity of the Balloon Analog Risk Task (BART): Associations With Psychopathy and Impulsivity. Assessment 12, 416–428 (2005).

26. White, T. L., Lejuez, C. W. & de Wit, H. Test-retest characteristics of the Balloon Analogue Risk Task (BART). Exp. Clin. Psychopharmacol. 16, 565–570 (2008).

27. Pearson, J. M., Hickey, P. T., Lad, S. P., Platt, M. L. & Turner, D. A. Local Fields in Human Subthalamic Nucleus Track the Lead-up to Impulsive Choices. Front. Neurosci. 11, (2017).

28. Clark, W. A. & Farley, B. G. Generalization of pattern recognition in a self-organizing system. in Proceedings of the March 1-3, 1955, western joint computer conference on - AFIPS’55 (Western) 86– 91 (ACM Press, Los Angeles, California, 1955). doi:10.1145/1455292.1455309.

29. Barto, A. G., Sutton, R. S. & Anderson, C. W. Neuronlike adaptive elements that can solve difficult learning control problems. IEEE Trans. Syst. Man Cybern. SMC-13, 834–846 (1983).

30. Schonberg, T., Fox, C. R. & Poldrack, R. A. Mind the gap: bridging economic and naturalistic risk-taking with cognitive neuroscience. Trends Cogn. Sci. 15, 11–19 (2011).

31. Fascianelli, V. et al. Neural representational geometries reflect behavioral differences in monkeys and recurrent neural networks. Nat. Commun. 15, 6479 (2024).

32. Musall, S., Urai, A. E., Sussillo, D. & Churchland, A. K. Harnessing behavioral diversity to understand neural computations for cognition. Curr. Opin. Neurobiol. 58, 229–238 (2019).

33. Bowler, J. C., Azhar, D., Jensen, C. M., Lee, H.-W. & Heys, J. G. Structured experience shapes strategy learning and neural dynamics in the medial entorhinal cortex.

34. Mante, V., Sussillo, D., Shenoy, K. V. & Newsome, W. T. Context-dependent computation by recurrent dynamics in prefrontal cortex. Nature 503, 78–84 (2013).

35. Zoltowski, D., Pillow, J. & Linderman, S. A general recurrent state space framework for modeling neural dynamics during decision-making. in Proceedings of the 37th International Conference on Machine Learning 11680–11691 (PMLR, 2020).

36. Sadtler, P. T. et al. Neural constraints on learning. Nature 512, 423–426 (2014).

37. Huys, Q. J. M., Cruickshank, A. & Seriès, P. Reward-Based Learning, Model-Based and Model-Free. in Encyclopedia of Computational Neuroscience (eds Jaeger, D. & Jung, R.) 1–10 (Springer New York, New York, NY, 2014). doi:10.1007/978-1-4614-7320-6_674-1.

38. Lucantonio, F., Caprioli, D. & Schoenbaum, G. Transition from ‘model-based’ to ‘model-free’ behavioral control in addiction: Involvement of the orbitofrontal cortex and dorsolateral striatum. Neuropharmacology 76, 407–415 (2014).

39. Russek, E. M., Momennejad, I., Botvinick, M. M., Gershman, S. J. & Daw, N. D. Predictive representations can link model-based reinforcement learning to model-free mechanisms. PLOS Comput. Biol. 13, e1005768 (2017).

40. Sebold, M. et al. Model-Based and Model-Free Decisions in Alcohol Dependence. Neuropsychobiology 70, 122–131 (2014).

41. Groman, S. M., Massi, B., Mathias, S. R., Lee, D. & Taylor, J. R. Model-Free and Model-Based Influences in Addiction-Related Behaviors. Biol. Psychiatry 85, 936–945 (2019).

42. Schulman, J., Wolski, F., Dhariwal, P., Radford, A. & Klimov, O. Proximal Policy Optimization Algorithms. Preprint at 10.48550/arXiv.1707.06347 (2017).

43. Andrychowicz, M., et al. What Matters In On-Policy Reinforcement Learning? A Large-Scale Empirical Study. Preprint at 10.48550/arXiv.2006.05990 (2020).

44. Misra, A. et al. Methods for implantation of micro-wire bundles and optimization of single/multi-unit recordings from human mesial temporal lobe. J. Neural Eng. 11, 026013 (2014).

45. Brainard, D. H. The Psychophysics Toolbox. Spat. Vis. 10, 433–436 (1997).

46. Pelli, D. G. The VideoToolbox software for visual psychophysics: transforming numbers into movies. Spat. Vis. 10, 437–442 (1997).

47. Shoham, S. Robust clustering by deterministic agglomeration EM of mixtures of multivariate t-distributions. Pattern Recognit. 35, 1127–1142 (2002).

48. Davis, T. S. et al. LeGUI: A Fast and Accurate Graphical User Interface for Automated Detection and Anatomical Localization of Intracranial Electrodes. Front. Neurosci. 15, (2021).

49. Pedregosa, F. et al. Scikit-learn: Machine Learning in Python. Mach. Learn. PYTHON.

50. Paszke, A., et al. PyTorch: An Imperative Style, High-Performance Deep Learning Library. Preprint at 10.48550/arXiv.1912.01703 (2019).

